# Will an outbreak exceed available resources for control? Estimating the risk from invading pathogens using practical definitions of a severe epidemic

**DOI:** 10.1101/768853

**Authors:** R.N. Thompson, C.A. Gilligan, N.J. Cunniffe

## Abstract

Forecasting whether or not initial reports of disease will be followed by a severe epidemic is an important component of disease management. Standard epidemic risk estimates involve assuming that infections occur according to a branching process and correspond to the probability that the outbreak persists beyond the initial stochastic phase. However, an alternative assessment is to predict whether or not initial cases will lead to a severe epidemic in which available control resources are exceeded. We show how this risk can be estimated by considering three practically relevant potential definitions of a severe epidemic; namely, an outbreak in which: i) a large number of hosts are infected simultaneously; ii) a large total number of infections occur; and iii) the pathogen remains in the population for a long period. We show that the probability of a severe epidemic under these definitions often coincides with the standard branching process estimate for the major epidemic probability. However, these practically relevant risk assessments can also be different from the major epidemic probability, as well as from each other. This holds in different epidemiological systems, highlighting that careful consideration of what constitutes a severe epidemic in an ongoing outbreak is vital for accurate risk quantification.

## 1. INTRODUCTION

Infectious disease epidemics in populations of humans, animals and plants represent a recurring risk worldwide [1–7]. An important aim for policy-makers near the start of an outbreak is to assess the risk posed by the invading pathogen, including whether initial cases will lead on to a major epidemic or whether the pathogen will die out rapidly instead [8,9]. An important practical consequence is that, if an outbreak is likely simply to fade out, then costly interventions such as vaccination [10,11], culling/felling/roguing of plants or agricultural animals [12–18] and workplace or school closure [19] may be unnecessary [20].

There is a well-known estimate for the probability of a major epidemic when a pathogen is newly arrived in a host population, which in its simplest form is given by

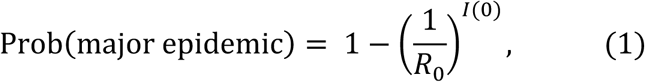

in which *R*_0_ is the basic reproduction number of the pathogen and *I*(0) is the number of individuals that are currently infected. The estimate in equation (1) applies to a wide range of models, including the commonly used Susceptible-Infected-Susceptible (SIS) and Susceptible-Infected-Removed (SIR) models [21]. It is derived by assuming that infections occur according to a branching process (see Methods). For the commonly used Susceptible-Exposed-Infectious-Removed model, the exponent in equation (1) would change from *I*(0) to *E*(0) + *I*(0) [9]. More sophisticated estimates that are based on similar branching process approximations can be derived for models including additional epidemiological detail, such as more complex population structure [22–24] and/or infectious periods that are not exponentially distributed [25,26].

The quantity in equation (1), and particularly the version in which *I*(0) = 1, is used extensively in the epidemiological modelling literature [8,9,21,26–35]. It is increasingly used in real-time during emerging outbreaks. For example, it was used during the 2014-16 epidemic of Ebola virus disease in West Africa to estimate the chance that, if the virus arrived in Nigeria, sustained transmission would follow in that country [30]. It was considered in the context of flare-ups in new locations for the recent Ebola epidemic in the Democratic Republic of the Congo [26]. Branching process models were also used at the start of the COVID-19 pandemic before cases were detected outside China to assess the risk of epidemics elsewhere [36,37], including the application of equation (1) [36].

However, while the major epidemic probability in equation (1) is useful to assess whether or not an outbreak is likely to persist beyond the initial stochastic phase, becoming a major epidemic does not guarantee that the outbreak will overwhelm available control resources. Over many outbreaks under identical conditions, if the population size is large and *R*_0_ is much greater than one, then the distribution of possible epidemic sizes is bimodal according to simple epidemic models such as the stochastic SIR model (Fig 1d – see also [38–42]). In other words, the final size of any single outbreak is almost always in one of two possible ranges. For example, in Fig 1d, virtually all outbreaks either lead to 1-20 hosts ever infected or to 700-860 hosts ever infected, where the precise ranges depend on the population size and the value of *R*_0_. The estimate for the probability of a major epidemic in equation (1) corresponds approximately to the proportion of outbreaks that have a final size in the higher of these ranges. The outbreaks within the higher range, however, do not necessarily represent outbreaks in which available control resources are exceeded. For practical assessments of the threat from an invading pathogen, it would often therefore be appropriate for the notion of a severe epidemic to be grounded in consequences for disease control, depending on the specific system and outbreak under consideration.

**Figure 1.**
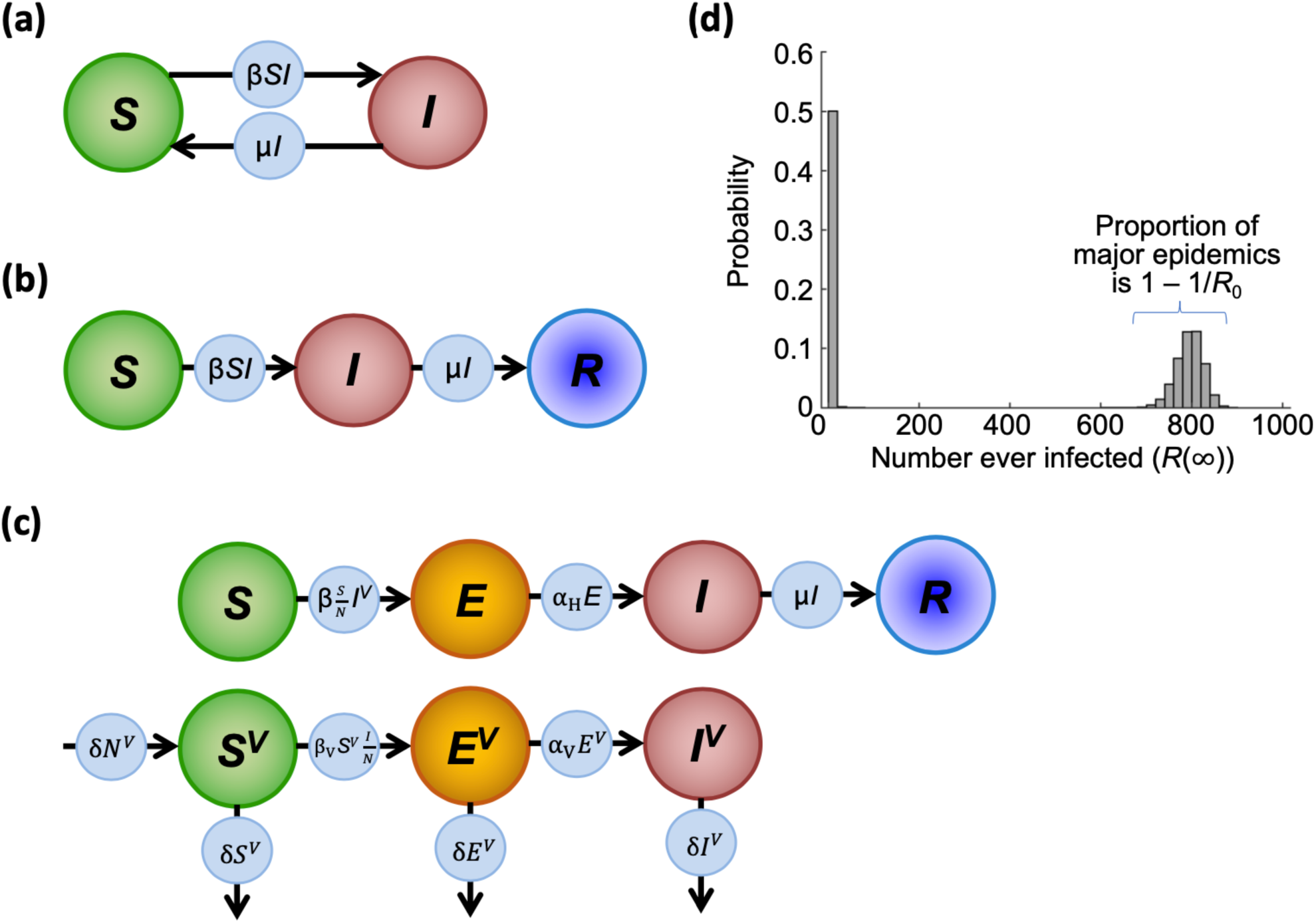
Schematic diagrams illustrating the population structures for the different models considered, and an example distribution of final sizes for the stochastic SIR model. (a) The SIS model. (b) The SIR model. (c) The host-vector model of Zika virus transmission. (d) Distribution of final sizes in the stochastic SIR model, with population size *N* = 1000, *R*_0_ = 2, *I*(0) = 1 and the rest of the population susceptible initially. The x-axis has been split into bars of width 20 (so that, for example, the first bar corresponds to the probability that between 1 and 20 individuals are ever infected).

Here, we assess whether or not outbreaks are likely to develop into severe epidemics according to three possible metrics that might be practically relevant in different outbreak scenarios. Specifically, these are:

### Concurrent size

In this assessment, a severe epidemic is an outbreak in which the number of individuals infected simultaneously exceeds the capacity for treatment.

### Total infections

In this assessment, a severe epidemic is an outbreak in which the total number of infections exceeds the number of available treatments.

### Duration

In this assessment, a severe epidemic is an outbreak that is not contained quickly and therefore persists for an unacceptably long period.

We compare the probability of a severe epidemic under each of these definitions, as well as calculate the branching process estimate for probability of a major epidemic (hereafter, we differentiate between the “probability of a severe epidemic” calculated using one of the metrics above and the “probability of a major epidemic” calculated by assuming that infections occur according to a branching process). In our analyses, as examples we consider three stochastic epidemiological models that are representative of different host responses to infection and capture different routes of transmission. Specifically, we consider the SIS model, the SIR model, and a host-vector model parameterised for Zika virus transmission. For the SIS and SIR models, the probability of a major epidemic corresponds to equation (1), and in the case of Zika virus the probability of a major epidemic is given by an adapted version of equation (1) that accounts for transmission between hosts by vectors (we present an approach for deriving these well-known formulae in Methods).

To motivate our analyses, we note that estimation of the risk of the outbreak going on to have a large concurrent size, a large total number of infections or a long duration might be the appropriate risk assessment in different scenarios. For example, it might be natural to assume that, if the number of individuals infected at any time always remains below the capacity for treatment, then the outbreak is not severe since medical care is available for all individuals that require treatment. Indeed, in the ongoing COVID-19 pandemic, one of the main aims of interventions in the UK has been to ensure that the number of individuals requiring intensive care unit beds remains below the total number of beds available [43–45]. More generally, the threshold capacity might derive from the number of available beds in hospitals or treatment units [46,47], or the availability of care workers [48]. This motivates consideration of the “concurrent size” metric above.

However, assessing the outbreak risk based on numbers of hosts infected simultaneously will not always be appropriate. Policy-makers often have to make decisions concerning how much treatment to stockpile; if all cases must be treated, this corresponds to the total number of infections during the outbreak. For example, in response to growing awareness of the threat of a potential influenza pandemic, between 2006 and 2013 policy-makers in the UK stockpiled around 40 million units of antivirals at a cost of £424 million. This led to severe criticism when only 2.4 million units were needed, the majority of which were used during the 2009 H1N1 influenza pandemic [49]. Another possible risk assessment is therefore whether or not the total number of infections will exceed a critical value (the “total infections” metric above). This critical value might be set by the stock of available treatments for use during the outbreak.

Finally, we consider a third possible risk assessment (using the “duration” metric). In this scenario, we evaluate whether or not an outbreak is likely to persist for an unacceptably long period. An outbreak that fades out quickly may escape public attention. Even if an outbreak leads to a significant number of hosts infected, if it ends relatively quickly then it might not be considered a severe epidemic. For example, the first Ebola outbreak in the Democratic Republic of the Congo in 2018 resulted in 53 cases, but was not considered a severe epidemic due to its fast containment, leading to commendation of the success of public health measures [50]. Consequently, an outbreak might only be classified as a severe epidemic if it persists for a threshold length of time.

An outbreak might be classified as a severe epidemic according to one of the metrics above, yet not be a severe epidemic if another metric is used. In 1665-66, plague affected the village of Eyam in the UK, which famously isolated itself via a self-imposed quarantine [51,52]. The outbreak in the village was long-running, and a large number of individuals were killed (most reports suggest 250-260 out of a total of 350 in the village died, although there is some uncertainty particularly regarding the size of the at-risk population [53]). However, model fits suggest that a maximum of only around 30 people were ever infected simultaneously [54–56]. As a result, this epidemic might have been classified as severe according to the “total infections” and “duration” metrics, yet not the “concurrent size” metric, depending on the precise values of the thresholds set in each case. This highlights the need to consider the appropriate metric for defining severe epidemics in the particular ongoing outbreak under consideration in order to perform the most practically relevant risk assessment.

A large body of theoretical work exists relating to the metrics for defining severe epidemics that we consider. For example, for the stochastic SIS and SIR models, probability distributions for the maximum number of individuals infected concurrently prior to epidemic extinction have been derived previously [57–59]. For *R*_0_ significantly greater than one, a related quantity (the quasi-stationary distribution – the distribution of the number of infected individuals in the long phase of the epidemic prior to extinction) has been studied in detail for models in which the pathogen persists long-term, including the stochastic SIS model [60–65] and the stochastic SIR model with births and deaths [66]. Analytic expressions and approximations have been found for the total number of infections over the course of an outbreak for a range of epidemic models [25,41,59,67– 69] and different approaches exist for calculating probability mass functions describing the possible final sizes of a stochastic epidemic (for a review of approaches for the stochastic SIR model, see [59]). The duration of an epidemic has also been well-studied [60,70–75], as has the duration of the initial stochastic phase of outbreaks that go on to become major epidemics [76].

However, previously developed mathematical theory is not the focus of our analyses. Instead, the novelty of the research that we present is to compare assessments of the risk from invading pathogens evaluated in different ways. We demonstrate the general principle that the precise definition of a severe epidemic (i.e. the metric chosen to define a severe epidemic, or the choice to use the standard branching process estimate for the major epidemic probability) can affect the assessments of the risk whenever a pathogen arrives in a new host population. The probabilities that an outbreak has a large concurrent size, a large total number of infections or a long duration may not coincide and depend on precisely which values of the relevant thresholds are set. These probabilities may or may not match the major epidemic probability assessed in the standard way. Careful consideration of precisely how a severe epidemic is classified is therefore necessary whenever the risk from an invading pathogen is estimated at the beginning of an emerging outbreak. Only once the notion of a severe epidemic has been formally defined – based on criteria of practical relevance – can this risk be properly assessed.

## 2. METHODS

We conduct five analyses. In the first three analyses, we consider the stochastic SIS model, the SIR model and the Zika host-vector model, and assess the risk that a single initial case will lead to an outbreak with a large “concurrent size”. Our final two analyses focus on the stochastic SIS model. We calculate the probability of an outbreak going on to exceed a pre-specified total number of infections (the “total infections” metric) or time (the “duration” metric).

Here, we describe the epidemiological models that we use, calculation of the branching process estimate of the major epidemic probability for each model, and how the probability of a severe epidemic can be calculated under the “concurrent size” metric for each of the models considered. We then explain how the probability of a severe epidemic under the other practically relevant metrics can be obtained for the SIS model, although our methodology generalises immediately to any model for which a method of stochastic simulation is available.

### Epidemiological models

#### Susceptible-Infected-Susceptible (SIS) model

According to the SIS model, at any time each individual in the population is classified to be either (*S*)usceptible to or (*I*)nfected by the pathogen. The deterministic SIS model is given by

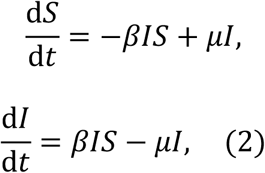

where *β* represents the infection rate between each susceptible-infected pair and *μ* is the rate at which each infected host recovers and becomes susceptible again. We use the analogous stochastic model in most of our analyses, in which the net rate at which any epidemiological event occurs is *βIS* + *μI*. At any time prior to the end of the outbreak, the probability that this next event is an infection is 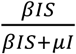 and the probability that the next event is a recovery is 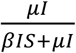.

In this model, if the total population size is *N*, the basic reproduction number is given by

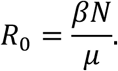

#### Susceptible-Infected-Removed (SIR) model

Under the SIR model, at any time each individual in the population is classified according to whether they are (*S*)usceptible to infection, (*I*)nfected, or (*R*)emoved and no longer spreading the pathogen or available for infection. The deterministic SIR model is given by

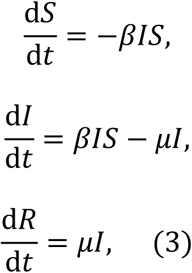

in which *β* is again a parameter governing the infection rate, and *μ* is the removal rate. In the analogous stochastic model, the net rate at which any epidemiological event occurs is still *βIS* + *μI*, and the probability that the next event is an infection event is similarly unchanged at 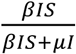. However, the other possible next event is a removal, which occurs with probability 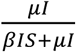. The basic reproduction number is again

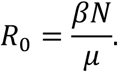

#### Zika transmission model

We consider the transmission of Zika virus according to a host-vector model [77], which we chose to demonstrate how the probability of a major epidemic can be calculated in a relatively complex epidemiological setting. In the model, the numbers of the *N* hosts that are (*S*)usceptible, (*E*)xposed, (*I*)nfectious and (*R*)emoved are tracked, as well as the number of the *N*^*V*^ vectors that are (*S*^V^)usceptible, (*E*^V^)xposed and (*I*^V^)nfectious. We adapt the version of the model as presented by Kucharski *et al*. [77] slightly to a more standard formulation in which all transmission terms are proportional to the relevant number of vectors and density of hosts (i.e., when most hosts are susceptible early in the outbreak, hosts are sufficiently abundant that the number of hosts that each vector bites is independent of the total number of hosts). The deterministic version of this model is then

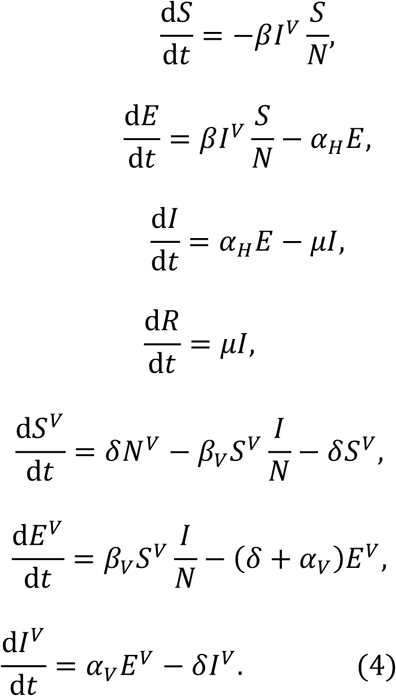

The parameters *β* and *β*_*V*_ govern the rates at which infectious vectors infect susceptible hosts and infectious hosts infect susceptible vectors, respectively. The mean latent periods of infections in hosts and vectors are given by 1/*α*_*H*_ and 1/*α*_*V*_. The parameter *μ* is the rate of removal of infectious hosts, and *δ* describes the death rate of every vector. In the analogous stochastic model, the number of infected human hosts arising from a single infected human (accounting for human-vector-human transmission) in an otherwise entirely susceptible population of humans and vectors is given by

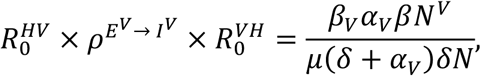

where 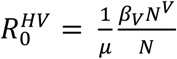 is the expected number of vectors infected (and going on to enter the exposed class) by a single infectious human, 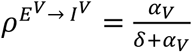 is the proportion of exposed vectors that become infectious and 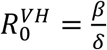 is the expected number of humans infected by a single infectious vector.

The basic reproduction number is given by

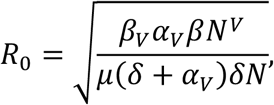

where the square root accounts for the fact that it takes two generations for infected humans to generate new infections, since new infections require host-vector-host transmission [78,79]. We note that in some studies, e.g. [77], the square root is omitted from the definition of *R*_0_. In contrast to the expression calculated by Kucharski *et al*. [77], to facilitate simulation of the stochastic model we explicitly track the total number of vectors, *N*^*V*^, rather than the density.

### Probability of a major epidemic (branching process estimate)

#### Standard estimate (stochastic SIS/SIR models)

The commonly used estimate for the major epidemic probability when a pathogen first arrives in a host population [8,9,35,36,21,27,29–34] can be derived by assuming that infections occur according to a branching process, making the assumptions that the susceptible population is large and that infection lineages arising from different infected hosts are independent. When a single infected host arrives in an otherwise susceptible population, the branching process estimate for the major epidemic probability is given by

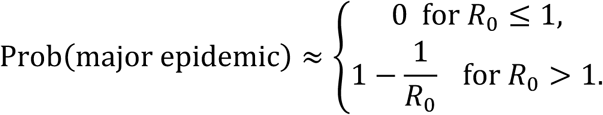

This expression is derived in Text S1.

If instead there are *I*(0) infected individuals initially rather than one, then for no major epidemic to occur, it is necessary for each initial infection lineage to die out, leading to the approximation given in equation (1) whenever *R*_0_ > 1.

#### Standard estimate (more complex models)

Here we show how the branching process estimate for the major epidemic probability can be derived for more complex systems by considering the stochastic version of the host-vector model given by the system of equations (4). Invasion probabilities have been estimated previously for vector-borne pathogens, for example by considering the distributions of secondary infections from infected hosts/vectors and their associated probability generating functions [22], however we use an alternative approach here (a “first step analysis”, as in [80]) that is straightforward to understand and extend to a range of epidemiological settings.

We denote the probability of no major epidemic occurring starting from *i* exposed or infectious human hosts, *j* exposed vectors and *k* infectious vectors by *q*_ijk_. We must consider exposed and infectious vectors separately to account for the possibility that exposed vectors die before becoming infectious.

Starting from a single infectious host introduced into an entirely susceptible population of hosts and vectors, and conditioning on the next event, gives

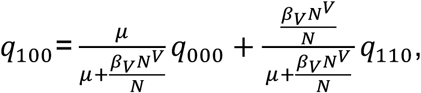

because the next event can then either be recovery of the infectious host (with probability 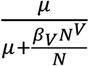) or infection of a susceptible vector (with probability 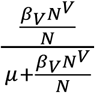). Similarly, starting from a single exposed or infectious vector gives

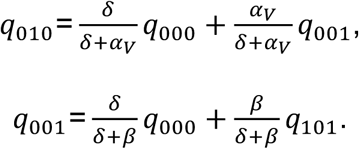

We again assume that infection lineages are independent, permitting us to approximate terms with two exposed or infectious individuals by non-linear terms involving single exposed or infectious individuals, e.g. *q*_110_ ≈ *q*_100_*q*_010_. Noting that *q*_000_ = 1, the three equations above can be solved to give expressions for *q*_100_, *q*_010_ and *q*_001_. In particular, the probability of a major epidemic starting from a single infected host is then

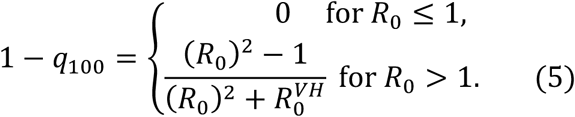

In this expression, 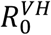 is the expected number of humans infected by a single infectious vector in an otherwise entirely susceptible population of humans and vectors.

### Probability of a severe epidemic (“concurrent size” metric)

As described in the introduction, we first define a major epidemic to be an outbreak in which the maximum number of individuals infected simultaneously is above a threshold value, which we denote by *M*. The value of *M* of relevance in practical applications might be set by the capacity for treatment.

#### Stochastic SIS model

Under the stochastic SIS model, the probability that the number of individuals infected simultaneously is at least *M* at some time prior to epidemic extinction can be calculated analytically [57]. This is advantageous since approximating this quantity using model simulations can be time consuming given that outbreaks under the SIS model can persist for long periods.

We denote the probability that the maximum number of individuals simultaneously infected (also known as the “peak prevalence” [59]) is at least *M*, starting from *I* infected individuals, as *q*_*I*_, and assume that the rest of the population is susceptible. Conditioning on whether the first event is an infection or recovery event gives

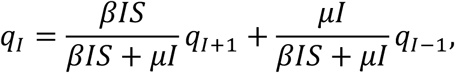

for *I* = 1,2,…,*M*-1. We solve this tridiagonal system of equations with boundary conditions *q*_0_ = 1 and *q*_M_ = 0, since the peak number infected will certainly be less than *M* if there are initially no infected individuals (assuming *M* > 0), and will be at least *M* if there are initially *M* infected individuals. Noting that *S* = *N* − *I*, these equations can be put in the form

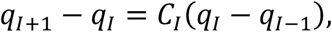

Where

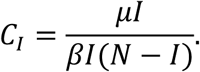

Iterating this gives

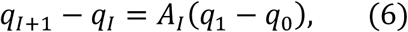

Where

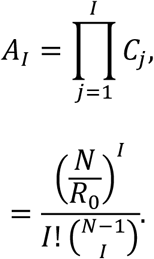

Adding equation (6) to itself for *I* = 1,2,…,*M* - 1 and rearranging gives

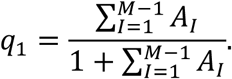

Instead adding equation (6) to itself for *I* = 1,2,…,*k* – 1 gives

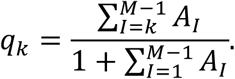

The probability of a severe epidemic according to the “concurrent size” metric is then given by

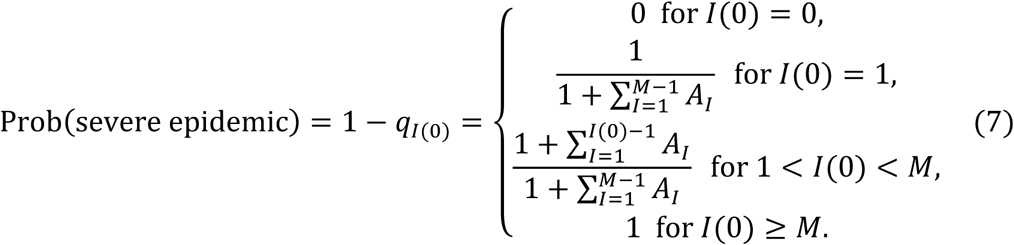

#### Stochastic SIR model

Under the stochastic SIR model, the probability that the maximum number infected simultaneously is at least *M* (i.e. a severe epidemic occurs, under the “concurrent size” metric) starting from any state (*I,R*) is calculated using an iterative approach [58,59]. Denoting the probability of a severe epidemic starting from state (*I,R*) by *p*_*I,R*_, then conditioning on the next event gives

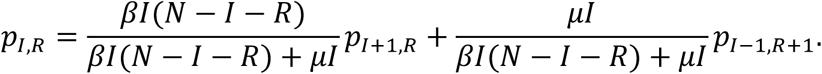

This system can be solved with boundary conditions *p*_0,*R*_ = 0, *p*_*I,N*−*M*+1_ = 0 and *p*_*M,R*_ = 1. Doing so does not require this system of equations to be solved simultaneously. Instead, the value of *p*_*I,R*_ is deduced for the following states (in order): (*I,R*) = (*M*-1,*N*-*M*), (*M*-2,*N*-*M*), …, (1,*N*-*M*), (*M*-1,*N*-*M-*1), …,(1,*N*-*M-*1),… (*M*-1,0),…,(1,0). For a schematic showing the order in which these probabilities are deduced, see Fig S1.

#### Zika transmission model

The probability of a severe epidemic using the “concurrent size” metric is approximated by simulating the model 10,000 times using the Gillespie direct method [81], and then calculating the proportion of simulations in which the number of infected human hosts is at least *M* at any point during the simulation.

### Probability of a severe epidemic (“total infections” and “duration” metrics)

We also consider the probability of a severe epidemic according to the stochastic SIS model for the “total infections” and “duration” metrics. Specifically, we estimate the probability that at least *F* infections occur over the course of the outbreak (prior to outbreak extinction), and the probability that the outbreak persists for at least *T* days.

We approximate these quantities by simulating the model 10,000 times using the Gillespie direct method [81] and recording separately the proportion of simulations in which there are at least *F* infections or that have a duration of at least *T* days. Each simulation is stopped when either of these following two criteria are satisfied: i) the simulated outbreak has gone extinct (*I* = 0), or; ii) both the number of infections has reached the maximum value of *F* considered (*F* = 2000; see Fig 5a) and the duration has reached the maximum value of *T* considered (*T* = 6000; see Fig 5b).

## 3. RESULTS

To begin to understand outbreak dynamics under the SIS, SIR and Zika transmission models, we first numerically solved the deterministic models given by the systems of equations (2), (3) and (4) with *R*_0_ = 1.5 in each case (Fig 2). For the parameter values considered, in a deterministic setting the SIS model predicts the largest number of individuals infected simultaneously as well as the most infections in total. Epidemics persisted forever (i.e. *I* remained larger than zero) under all three models, although the number of infected hosts tended to zero under the SIR and Zika transmission models.

**Figure 2.**
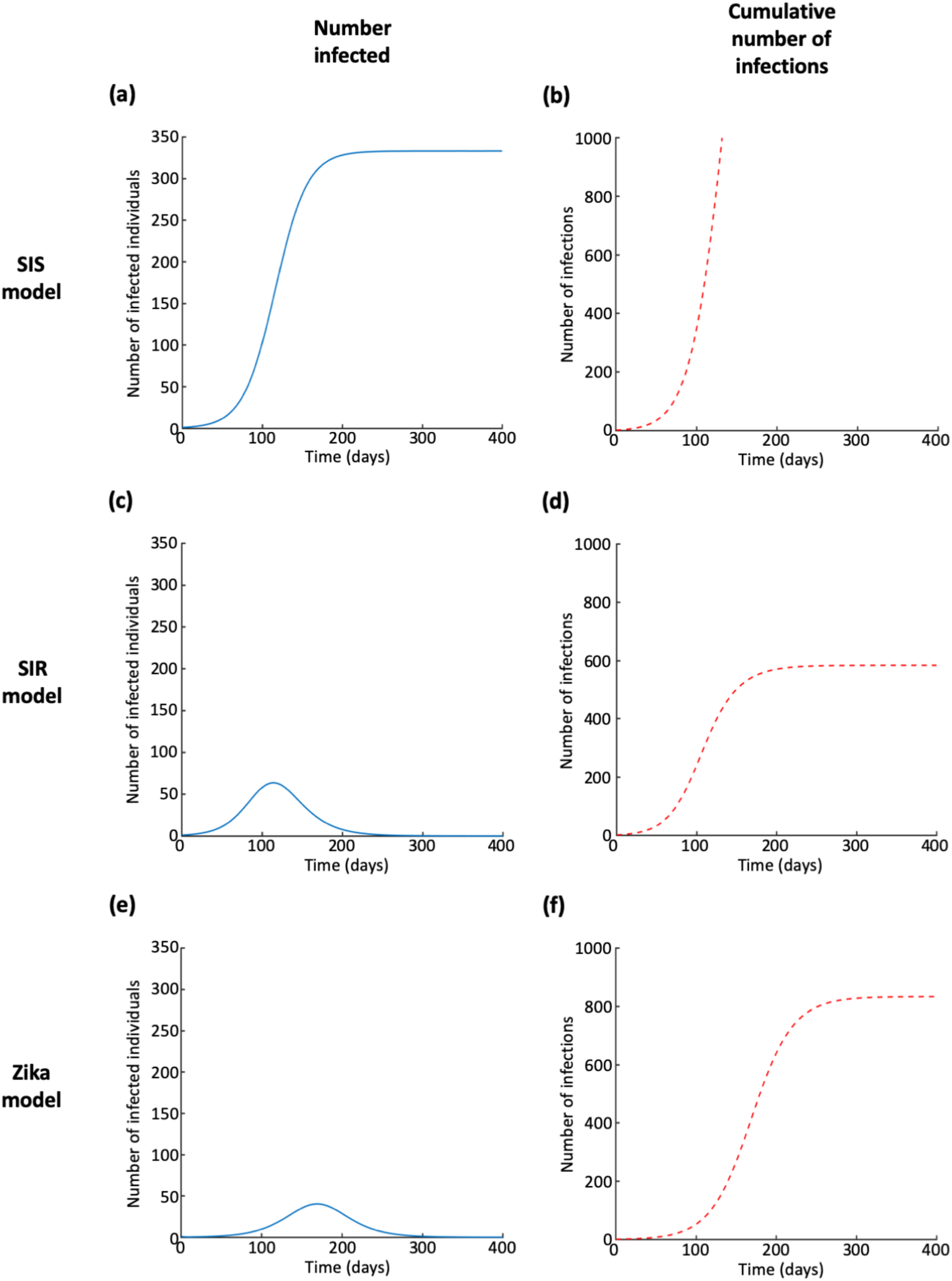
Numerical solutions of the deterministic SIS, SIR and Zika virus transmission models when the basic reproduction number is *R*_0_ = 1.5. (a) Temporal changes in the number of infected individuals according to the deterministic SIS model. (b) Temporal changes in the cumulative number of infections according to the deterministic SIS model. (c)-(d) Equivalent to a-b but for the deterministic SIR model. (e)-(f) Equivalent to a-b but for the deterministic Zika virus transmission model. Parameters for deterministic SIS and SIR models: *N* = 1,000, *β* = 0.00015 per day, 1/*μ* = 10 days. Parameters for deterministic Zika virus transmission model: *N* = 1,000, *N*^*V*^ = 10,000, 1/*α*_*V*_ = 10.5 days, 1/*α*_*H*_ = 5.9 days, 1/*μ* = 5 days, 1/*δ* = 7.8 days, *β* = 6.15 × 10^−2^ per day, *β*_*V*_ = 0.22 per day [77].

However, our main focus is assessing the risk from an invading pathogen, according to the “concurrent size”, “total infections” and “duration” metrics for defining severe epidemics. We therefore considered the more realistic stochastic models, in which demographic stochasticity is included. In the following sections, first we calculate the probability of a severe epidemic for the stochastic SIS model under the “concurrent size” metric. We then consider different epidemiological models, as well as other metrics defining a severe epidemic. In each case, the probability of a severe epidemic for the particular epidemiological model-severe epidemic metric pair under consideration is compared with the branching process approximation to the probability of a major epidemic for that model. The rationale for this comparison is that both quantities represent a possible way to assess the risk from an invading pathogen. Results are shown in Figs 3-5 and summarised in Tables 1 and 2. A chart outlining the model-severe epidemic metric pairs considered in each figure is shown in Fig S2.

**Figure 3.**
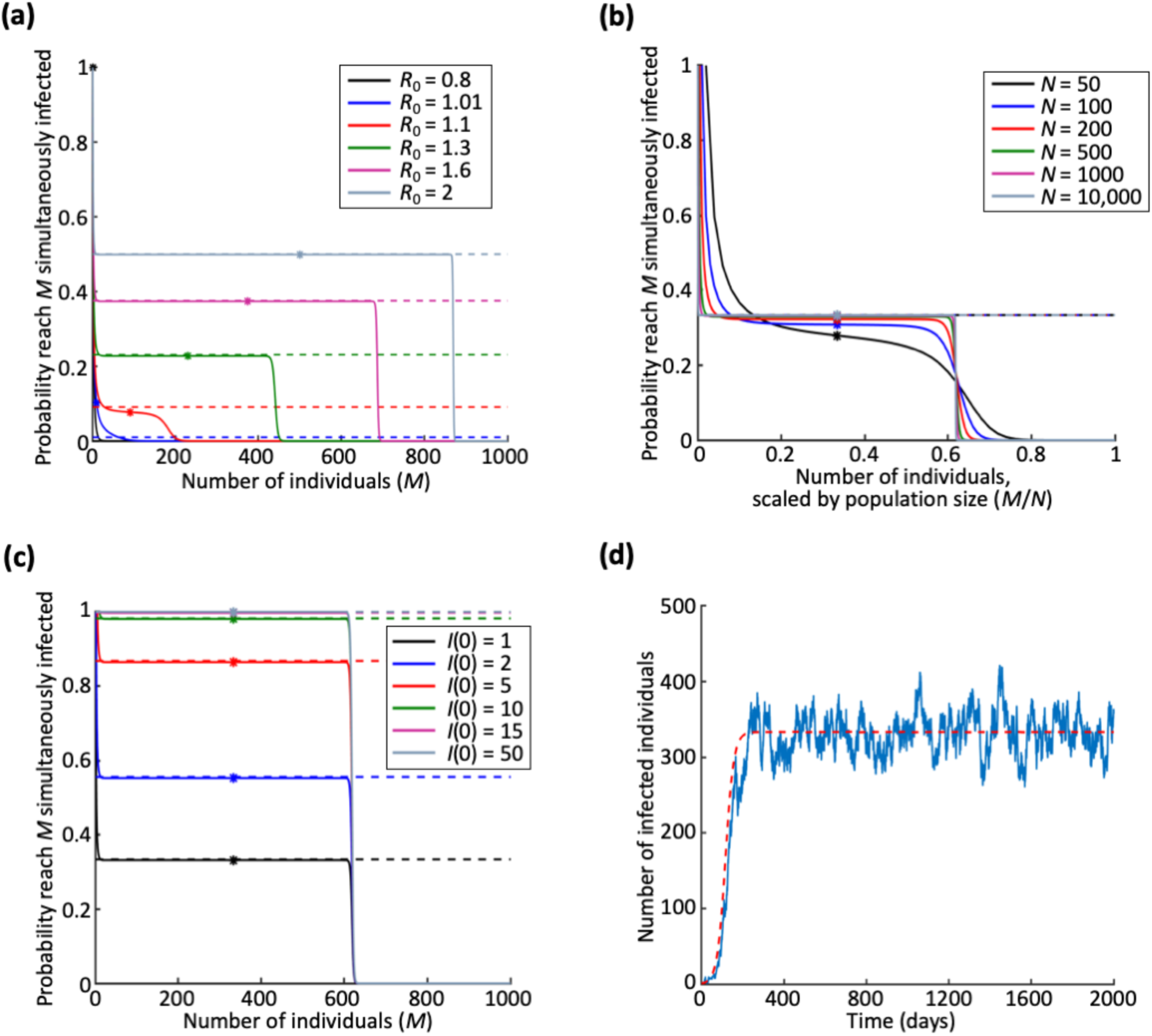
Probability of a severe epidemic under the SIS model, where a severe epidemic is defined as an outbreak in which at least *M* individuals are infected simultaneously at some time during the outbreak (“concurrent size” metric). (a) Dependence on *R*_0_. Solid lines represent the probability of a severe epidemic (system of equations (7)), dotted lines represent the branching process estimate for the major epidemic probability (equation (1)), and dots show the maximum number simultaneously infected in the analogous deterministic models (calculated analytically as shown in Text S1). *R*_0_ is varied by changing the value of *β*. (b) Equivalent to a, but showing dependence on the population size, *N*. (c) Equivalent to a, but showing dependence on the initial number of infected individuals, *I*(0). (d). Single simulation of the stochastic SIS model (blue), and numerical solution of deterministic SIS model (red dotted). The value of *I* in the stochastic simulation will continue to fluctuate about the deterministic value until *I* reaches 0. Parameter values (except where stated): *N* = 1,000, *R*_0_ = 1.5, *I*(0) = 1 and *R*(0) = 0 and the remainder of the population susceptible initially. In panel d, *β* = 0.000015 per day and 1/*μ* = 10 days.

**Table 1.** Effect of underlying epidemiology. For which values of the threshold (number of hosts simultaneously infected, *M*) in the “concurrent size” definition of a severe epidemic does the probability of a severe epidemic match the branching process estimate for the major epidemic probability? Range of values here are those for which the probability of a severe epidemic is within 0.01 of the probability of a major epidemic. Results are summarised for the SIS, SIR and Zika virus models, for the parameter values shown in the relevant figure captions.

|  | **SIS model** | **SIR model** | **Zika host-vector model** |
| --- | --- | --- | --- |
| $R_0 = 1.1$ | $21 \leq M \leq 54$<br>(Fig 3a) | $18 \leq M \leq 21$<br>(Fig 4a) | $6 \leq M \leq 7$<br>(Fig 4b) |
| $R_0 = 1.6$ | $8 \leq M \leq 681$<br>(Fig 3a) | $8 \leq M \leq 55$<br>(Fig 4a) | $3 \leq M \leq 47$<br>(Fig 4b) |
| $R_0 = 2$ | $6 \leq M \leq 866$<br>(Fig 3a) | $7 \leq M \leq 117$<br>(Fig 4a) | $2 \leq M \leq 83$<br>(Fig 4b) |

### The probability of a major epidemic

We calculated the probability of a severe epidemic according to the stochastic SIS model under the “concurrent size” metric for a severe epidemic – i.e. an outbreak in which the number of individuals infected simultaneously is at least a pre-specified threshold number (*M*) at some time during the outbreak. In this case, as described in Methods, it is possible to calculate the probability of a severe epidemic analytically.

We show the probability of a severe epidemic for a range of values of the threshold *M* in Fig 3a. Of course, for *R*_0_ much larger than one, the probability of a severe epidemic was approximated closely by the standard branching process estimate for the probability of a major epidemic for many values of the threshold, *M*. When, however, *R*_0_ was close to one, the standard estimate corresponded to a single choice of *M* (see e.g. blue lines in Fig 3a, where the solid line is close to the corresponding dotted line in only one place, i.e. for a single value of *M*). The parameter regime in which *R*_0_ is close to one is important in many real epidemiological systems since the aim of pre-emptive control strategies is often to reduce *R*_0_ below one (or, when an outbreak has started, to reduce the time-varying or effective reproduction number below one [82–85]).

In large host populations, the probability of a severe epidemic as a function of *M* took the form of a step function (Fig 3b). This suggested that, if the pathogen successfully invaded the population, then the number of infected individuals would definitely reach a specific maximum value which is determined by *R*_0_. For example, for outbreaks with *R*_0_ = 1.5, the pathogen will invade the population with probability 0.33 (i.e. the probability of a major epidemic), and, if this occurs, then around two-thirds of the population will be infected simultaneously at some time during the epidemic.

An approximation to the maximum value of *I* that will be reached in the stochastic SIS model in the large *N* case is given by

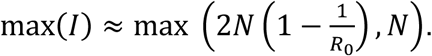

The expression 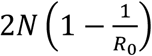 is twice the maximum value of *I* in the corresponding deterministic model. This approximation involves the assumption that, if the pathogen invades in the stochastic SIS model (so that a major epidemic occurs), *I* will fluctuate approximately symmetrically around the deterministic equilibrium value (Fig 3d). This is consistent with the previous finding that, in the stochastic SIS model, the quasi-stationary distribution can be approximated by the (symmetric) normal distribution when *R*_0_ is larger than and not close to one [62]. It is likely, then, that by the time the pathogen dies out by reaching *I* = 0, the number of infected individuals will at some stage also have reached approximately double the value it fluctuated around too. In Fig 3b, when *N* = 10,000, this approximation gives a maximum proportion of the population simultaneously infected of 0.67, when the true value is 0.62.

We also note that, if the expression above for max(*I*) is reformulated to give the maximum proportion of the population that is simultaneously infected, the resulting expression is independent of the population size *N*. This can also be seen graphically – in Fig 3b, if the pathogen successfully invaded the population then the maximum proportion of hosts that were infected simultaneously was approximately independent of the size of the host population, so long as *N* was sufficiently large.

### Different epidemiological models

We considered the probability that the maximum number of individuals infected simultaneously is at least a pre-specified threshold (i.e. a severe epidemic occurs, using the “concurrent size” metric) under the SIR and Zika virus transmission models. For the stochastic SIR model, we used an iterative method to calculate this probability as described in Methods.

For the stochastic Zika virus transmission model, we simulated the model in a population of *N* = 1,000 human hosts and *N*^V^ = 10,000 vectors using the Gillespie direct algorithm [81], using parameter values from Kucharski *et al*., 2016 [77] – see caption of Fig 4. The value of *R*_0_ was then varied in Fig 4b by altering the parameter *β* that governs the rate at which infected vectors infect susceptible human hosts.

**Figure 4.**
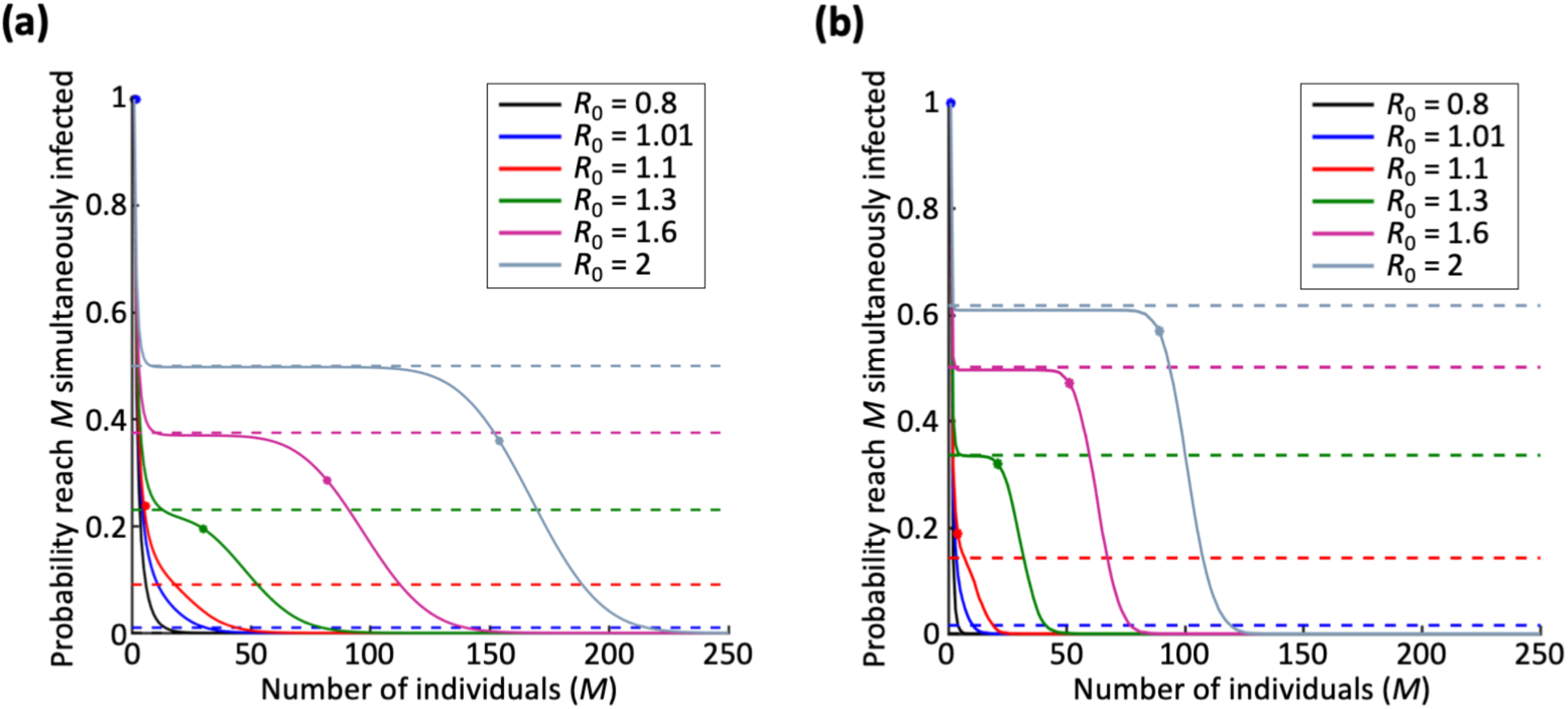
Probability of a severe epidemic under the SIR and Zika virus transmission models, where a severe epidemic is defined as an outbreak in which at least *M* individuals are infected simultaneously at some time during the outbreak (“concurrent size” metric). (a) SIR model. Solid lines represent the true probability of a severe epidemic calculated using the iterative method described in Methods, dotted lines represent the branching process estimate of the probability of a major epidemic (equation (1)), and dots show the maximum number simultaneously infected in the analogous deterministic model (calculated analytically for the SIR model as shown in Text S1). *R*_0_ is varied by changing the value of *β*. (b) Equivalent to a, but for the Zika virus transmission model (where *M* refers to the number of simultaneously infected hosts). For the Zika transmission model, the probability of a severe epidemic is calculated by simulation, and the branching process estimate of the major epidemic probability is given by equation (5). The maximum number simultaneously infected is found for the deterministic Zika virus transmission model by numerically solving the model. For both models, *N* = 1,000. Other parameters for the Zika virus transmission model: *N*^*V*^ = 10,000, 1/*α*_*V*_ = 10.5 days, 1/*α*_*H*_ = 5.9 days, 1/*μ* = 5 days, 1/*δ* = 7.8 days, *β*_*V*_ = 0.22 per day [77]. Initial conditions for both models comprise of a single infected host, with all other individuals (for the Zika transmission model, hosts and vectors) susceptible.

Under the stochastic SIR and Zika virus transmission models, for *R*_0_ larger than and not close to one, the maximum number of simultaneously infected individuals whenever the pathogen invaded the host population was typically smaller than under the SIS model (*cf*. Fig 2). Nonetheless, we found qualitatively similar behaviour in these cases – the probability of a severe epidemic was similar to the major epidemic probability approximated using a branching process for a wide range of values of the severe epidemic threshold when *R*_0_ was high (Fig 4). However, even if that is the case, the practically relevant value of the severe epidemic threshold (e.g. the number of available hospital beds) may mean that the severe epidemic probability does not match the major epidemic probability. For example, if *R*_0_ = 2 and 250 beds are available, for the SIR model the probability of a severe epidemic under the “concurrent size” definition is 0 (solid grey line in Fig 4a), yet the branching process estimate for the probability of a major epidemic is 0.5 (dotted grey line in Fig 4a).

### Alternative definitions of a severe epidemic

For the stochastic SIS model, we then calculated the probability of a severe epidemic using different metrics to define a severe epidemic – specifically, outbreaks in which there are at least *F* infection events (the “total infections” metric – Fig 5a) or outbreaks that persist for at least *T* days (the “duration” metric – Fig 5b).

**Figure 5.**
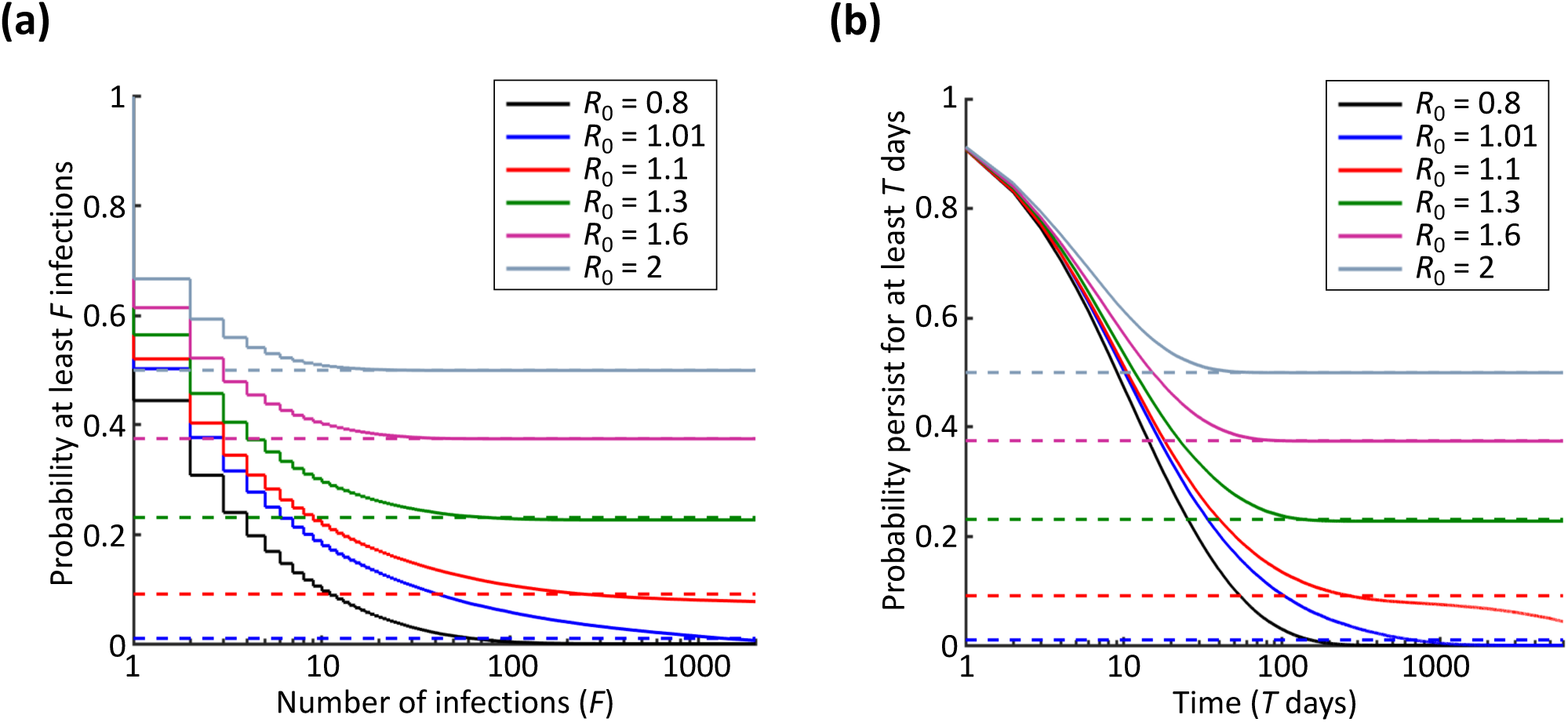
Probability of a severe epidemic under the SIS model, for different definitions of a severe epidemic. (a) A severe epidemic is defined as an outbreak in which at least *F* infections occur (“total infections” metric). (b) A severe epidemic is defined as an outbreak that persists for at least *T* days (“duration” metric). Solid lines represent the probability of a severe epidemic assessed via simulation of the stochastic model, and dotted lines represent the branching process estimate for the probability of a major epidemic (equation (1)). The x-axis is shown on a log-scale, for *F* between 1 and 2000 (panel a) and *T* between 1 and 6000 (panel b). The step function in panel a reflects the fact that the total number of infections can only take discrete values. Results of the deterministic model are not included in the figure, since under the deterministic SIS model epidemics persist indefinitely and generate an infinite number of infections whenever *R*_0_ > 1. Parameter values: *N* = 1,000, *I*(0) = 1 and *R*(0) = 0 and the remainder of the population susceptible initially. In both panels, *R*_0_ is varied by changing the value of *β*. In panel b, 1/*μ* = 10 days.

In the stochastic SIS model, if the pathogen invaded the host population then it tended to persist for long periods. Consequently, the probability of a severe epidemic using the “total infections” or “duration” metrics is approximately equal to the major epidemic probability for a wide range of values of the severe epidemic thresholds (i.e. values of *F* or *T*) compared with under the “concurrent size” definition. However, even in these cases, for small or very large values of the severe epidemic thresholds, the probability of a severe epidemic does not match the major epidemic probability, particularly when *R*_0_ is larger than but close to one (see e.g. red line in Fig 5b).

## 4. DISCUSSION

Evaluating the risk from an invading pathogen early in a potential severe epidemic is vital for planning interventions and determining whether or not current control resources are sufficient. When a pathogen arrives in a new location, the probability that initial cases will lead to a major epidemic as opposed to fading out as a minor outbreak can be approximated by assuming that infections occur according to a branching process. This probability represents the risk that the outbreak will persist beyond the initial stochastic phase in which case numbers are low. For simple models such as the stochastic SIS and SIR models, this corresponds to the major epidemic probability in equation (1). This can be extended to estimate the major epidemic probability using models with additional complexity, as we demonstrated by considering the case of host-vector transmission (see equations (4) and (5)).

However, the threat from an invading pathogen can also be assessed by estimating the probability that the outbreak will become “severe” and overwhelm available control resources. For example, the probability that the peak prevalence exceeds the capacity of treatment facilities (e.g. the number of hospital beds) can be estimated (Figs 3 and 4), as well as the probability that the outbreak exceeds a threshold in the total number of infections or has a long duration (Fig 5). In each case, the probability of a severe epidemic depends on the value of the threshold set (e.g. the exact number of hospital beds) to differentiate severe epidemics from other outbreaks, as well as the metric used to define a severe epidemic. This highlights the need to consider the precise definition of a “severe epidemic” carefully when assessing the risk from an invading pathogen.

When *R*_0_ is much greater than one or when the population size is extremely large, however, the probability of a severe epidemic is constant for a range of values of the threshold differentiating severe epidemics from other outbreaks (see e.g. different values of *M* in Fig 3a,b). In these cases, the probability of a severe epidemic will often match between definitions (Table 2). This is perhaps unsurprising since, for example, an outbreak with a large total number of infections may well also have a large peak prevalence. The probability of a severe epidemic is then approximately equal to the major epidemic probability calculated in the standard way (Table 2). However, even in this case, the specific threshold of practical importance in the outbreak may correspond to a different probability of a severe epidemic compared with the probability corresponding to a wide range of threshold values. Consequently, if a policy-maker wishes to understand the risk that an invading pathogen will overwhelm available resources for control, then this question should be addressed directly by choosing the relevant metric and threshold value carefully.

**Table 2.**
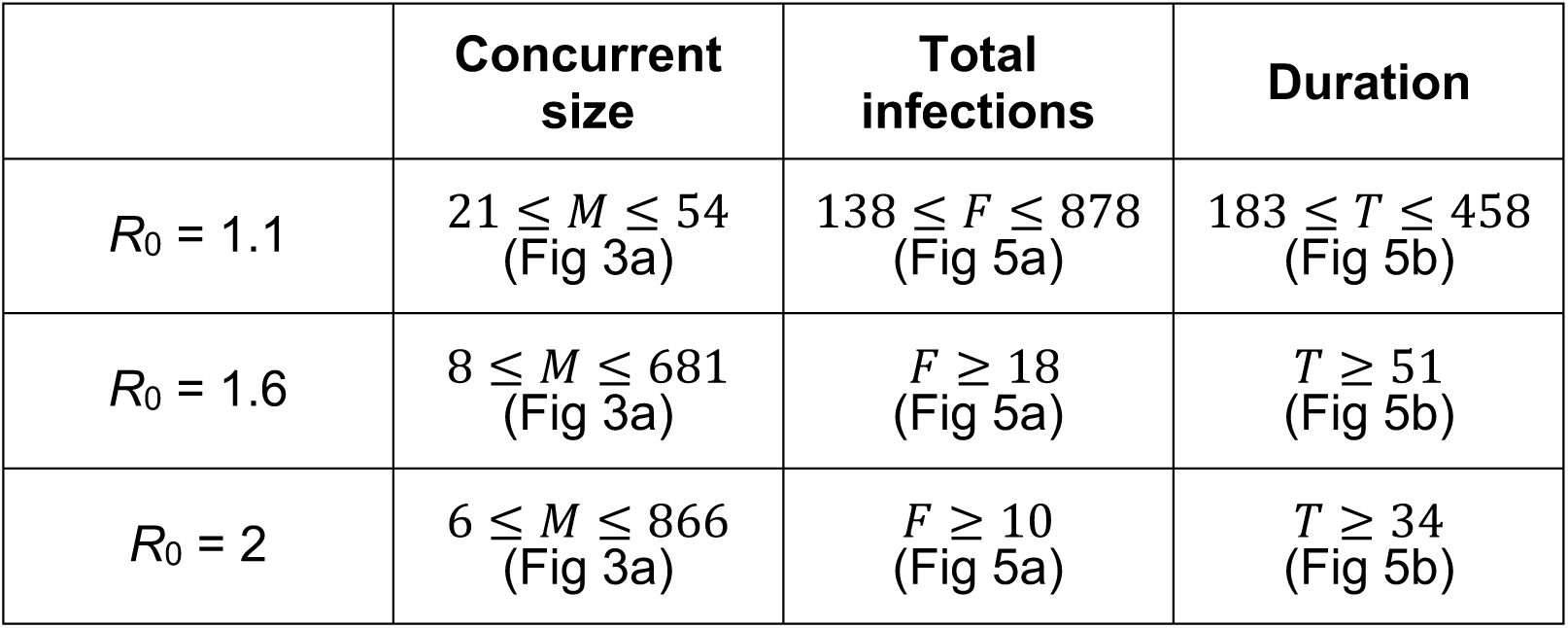
Effect of metric used to assess the probability of a severe epidemic. For which values of the thresholds in the practically relevant definitions of a severe epidemic do the probability of a severe epidemic match the branching process estimate for the major epidemic probability? Range of values here are those for which the probability of a severe epidemic is within 0.01 of the probability of a major epidemic. Results are summarised for the SIS model, for the parameter values shown in the relevant figure captions. “total infections” threshold values were tested up to a maximum of *F* = 2,000 (Fig 5a) and “duration” threshold values were tested up to *T* = 6,000 (Fig 5b).

|  | **Concurrent size** | **Total infections** | **Duration** |
| --- | --- | --- | --- |
| $R_0 = 1.1$ | $21 \leq M \leq 54$<br>(Fig 3a) | $138 \leq F \leq 878$<br>(Fig 5a) | $183 \leq T \leq 458$<br>(Fig 5b) |
| $R_0 = 1.6$ | $8 \leq M \leq 681$<br>(Fig 3a) | $F \geq 18$<br>(Fig 5a) | $T \geq 51$<br>(Fig 5b) |
| $R_0 = 2$ | $6 \leq M \leq 866$<br>(Fig 3a) | $F \geq 10$<br>(Fig 5a) | $T \geq 34$<br>(Fig 5b) |

We considered practical definitions of a severe epidemic that were based on thresholds such as the availability of treatment. A previous study defined severe epidemics according to a threshold in the percentage of the population ever infected, and concluded that epidemiological modellers should report the precise cutoff used to define such epidemics in model simulations [86]. Their conclusion was based on the observation that different thresholds in the percentage of hosts ever infected corresponded to wide variations in the other outputs of model simulations including the number of dead hosts or the time of the epidemic peak. We support this conclusion, and indeed some authors have reported the definition of a severe epidemic they used clearly – for example, Keeling *et al*. [87] differentiate between outbreaks in which less than one-third of the population becomes infected and those in which larger numbers of individuals are ever infected. Similarly, a recent study explored the risk of resurgence of COVID-19 when interventions are removed in different countries [88]. In that study, resurgence was said to have occurred when the number of individuals who are symptomatic infectious reached 100. However, while these values were reported clearly, we emphasise that the precise type of threshold and the value used should be chosen according to practical relevance in the particular system under consideration.

Under the first definition of a severe epidemic that we considered (using the “concurrent size” metric), the probability of a severe epidemic was assessed in the context of the capacity for treatment by estimating whether or not a threshold number of simultaneously infected individuals was likely to be exceeded. This definition may be practically relevant in a range of scenarios. For example, real-time analysis of a diphtheria epidemic in Cox Bazar’s in Bangladesh involved assessing the number of hospital beds that were needed [46]. The number of beds required was approximated in that study by using a model to forecast disease incidence, assuming that 15% of reported cases would require treatment as inpatients with an average hospital stay of five days for each case. The number of hospital beds that were already available might have provided a practically relevant severe epidemic threshold. Another example for which this type of threshold might apply is citrus greening disease in Brazil, where a law was introduced stating that a citrus grove must be destroyed if 28% of trees in the grove were infected and symptomatic [89,90]. At a local spatial scale, severe epidemics could therefore be defined as outbreaks in which more than 28% of trees in a grove are infected and symptomatic concurrently. Other examples for which interventions were introduced as soon as a threshold in the number simultaneously infected was reached include the development of the National Chlamydia Screening Programme in the United Kingdom in 2002 in response to the large size of the infected population [91].

However, no single metric for determining whether or not an outbreak is a severe epidemic will be relevant in all situations. To illustrate the ambiguity to which this might lead, we also considered two other definitions of a severe epidemic. In one of these (when the “total infections” metric was used), whether or not an outbreak was classified as a severe epidemic referred to the total number of infection events over the course of the outbreak, rather than the maximum number simultaneously infected. This might correspond to the total number of treatments required, which may be an important threshold if treatments have been stockpiled prior to the outbreak [49]. This definition might also be relevant if, for example, a policy-maker has to choose how to deploy resources between two different epidemics. If there are only sufficient resources to contain one outbreak, and both epidemics are equally controllable, then – in the absence of other considerations – it might be preferable to choose to contain the epidemic that is likely to generate more infections. In other real-world scenarios, alternative definitions might be appropriate. We also considered classifying severe epidemics as outbreaks that persist beyond a threshold length of time (using the “duration” metric). Different definitions of a severe epidemic might appear contradictory – for example, treatment can act to reduce the total number of infections yet increase the duration of the outbreak [92], making a severe epidemic less likely when the “total infections” metric is used but more likely when instead the “duration” metric is used.

Our intention here was to use very simple models to demonstrate the principle that different approaches for evaluating the threat from an invading pathogen can lead to very different assessments of risk. As described in the introduction, our research builds on a rich history of analyses that relate to the results obtained here. For example, for the stochastic SIS model, the time to extinction varies with *R*_0_. For *R*_0_ < 1, outbreaks will certainly end quickly. When *R*_0_ > 1, however, if *R*_0_ is increased then the duration of the epidemic also increases (exponentially with *R*_0_ [62]). Exact analytic expressions for the duration of a stochastic SIS epidemic have not been found, and so we assessed the probability of a severe epidemic under the “duration” metric using model simulations (Fig 5). However, analytic approximations to the expected duration of a major epidemic exist (e.g. [60,75]), and exploring the relationship between these approximations and the probability of a severe epidemic for different values of the threshold under the “duration” metric represents an interesting avenue for further investigation.

Another important extension of the research presented here is to explore the risk of a severe epidemic for outbreaks that do not assume that the population is well-mixed. The field of contact network epidemiology provides a framework in which the risk from an invading pathogen can be explored, accounting for the topology of the underlying network when making epidemiological predictions and planning public health measures. In that context, the probability of a major epidemic can be derived, depending on the transmissibility (*T*) of the pathogen rather than *R*_0_ [93]. The value of *T* represents the average probability that an infected host will transmit the pathogen to a susceptible individual that they have contact with. Meyers *et al*. [94] investigated the dependence of the major epidemic probability on the degree of the index case, as well as the major epidemic probability for different numbers of initial cases, in the context of SARS. Those authors, as well as Pourbohloul *et al*. [95], explored the effects of different interventions that reduce the numbers of contacts on quantities including the probability of a major epidemic. The framework underlying these models has also been extended to account for the time evolution of outbreaks [76], which is imported if the evaluation of the risk from an invading pathogen is to be linked to the extinction time of the outbreak (as in the “duration” metric that we considered).

Although our approach could be extended for different types of models (such as network models), compartmental models (such as the SIS and SIR models) are commonly used for assessing outbreak risks. Accurate outbreak forecasting using a compartmental model requires the model to be carefully matched to the epidemiology of the host-pathogen system, potentially including within-host dynamics [96,97], asymptomatic transmission [9,98,99] or spread between spatially distinct regions [29,100]. For certain definitions of a severe epidemic, it may be necessary to include bed-ridden or convalescent hosts in the model explicitly. For example, if the definition of a severe epidemic is linked to the availability of beds in treatment centres (as may be the case when the “concurrent size” metric is used), then infected individuals in treatment centres could be included in the model explicitly (for an example in which we consider three different models of an Ebola epidemic with different levels of complexity, see Text S3). More complex definitions of severe epidemics could be used, potentially requiring multiple criteria to be satisfied for an outbreak to be classified as a severe epidemic. In these more complicated scenarios, analytic calculations of the probability of a severe epidemic might not be possible. Model simulations can then be used to assess the risk from the invading pathogen, as we showed for a host-vector model of Zika virus transmission (Fig 4b).

We note that practical use of the methods presented here at the start of an emerging outbreak to assess the outbreak risk might require the parameters governing pathogen transmission to be estimated directly from case notification data. A range of methods exist for estimating reproduction numbers in real-time during outbreaks [83,84,101,102], including those designed for estimation in the early stochastic phase [103,104]. Practical use of the approaches we have developed might also require the wide range of interventions that are introduced in outbreak response settings to be integrated into the models explicitly. One way in which control can be included is to consider the effective reproduction number when the pathogen arrives in the system instead of the basic reproduction number, since the effective reproduction number accounts for interventions [26,82–84,105–107]. In that situation, the results that we presented would be unchanged (except that e.g. the lines in Fig 3a would correspond to different values of the effective reproduction number rather than the basic reproduction number). Temporal changes in interventions once an outbreak is underway have been approximated in epidemiological models by assuming that the values of the parameters governing transmission change during the outbreak, either by assuming that transmissibility changes at single timepoints [108–110] or continuously as the outbreak progresses [111,112]. However, for detailed descriptions of control to be included in estimates of the severe epidemic risk, more complex interventions should be included in model simulations. Since models are often used to test possible control strategies [7,12,20,45,82,113–115], this is a simple extension of the results presented here.

In summary, we have shown that the precise definition of a severe epidemic should be considered carefully in future studies that aim to evaluate the risk when a pathogen first arrives in a host population. Only once a “severe epidemic” has been defined precisely for the specific outbreak and setting under consideration can the probability of a severe epidemic occurring be properly assessed. Providing an explicit demonstration of the consequences of not considering the practically relevant definition in evaluating the risk is the key contribution of this paper.

## SUPPLEMENTARY MATERIAL

Text S1.

Text S2.

Figure S1. Schematic showing how the probability of a severe epidemic under the “concurrent size” definition can be deduced for the stochastic SIR model. The probability corresponding to each state (*I,R*) is deduced iteratively in the order 1,2,3… (red). The black arrows indicate which previous values are used to inform each new deduction.

Figure S2. Chart indicating the model-severe epidemic metric pair considered in the figures in the main text.

## AUTHORS’ CONTRIBUTIONS

All authors designed the study. RNT undertook the research and wrote the first draft of the manuscript. All authors revised the manuscript and approved the final version for submission.

