## Supplementary figures and images for "Will an outbreak exceed available resources for control? Estimating the risk from invading pathogens using practical definitions of a severe epidemic"

### Fig S1

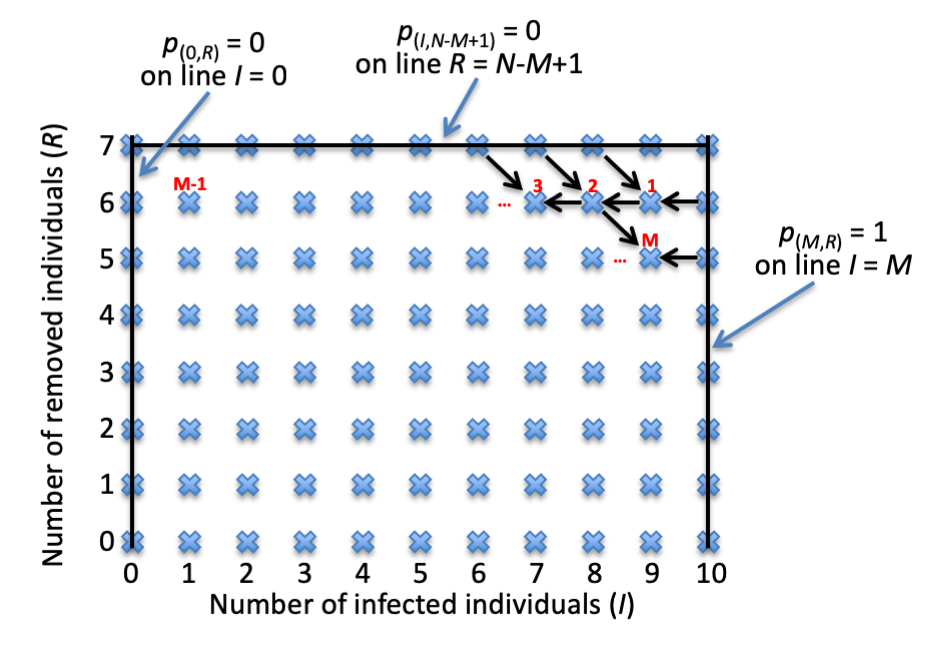

### Fig S2

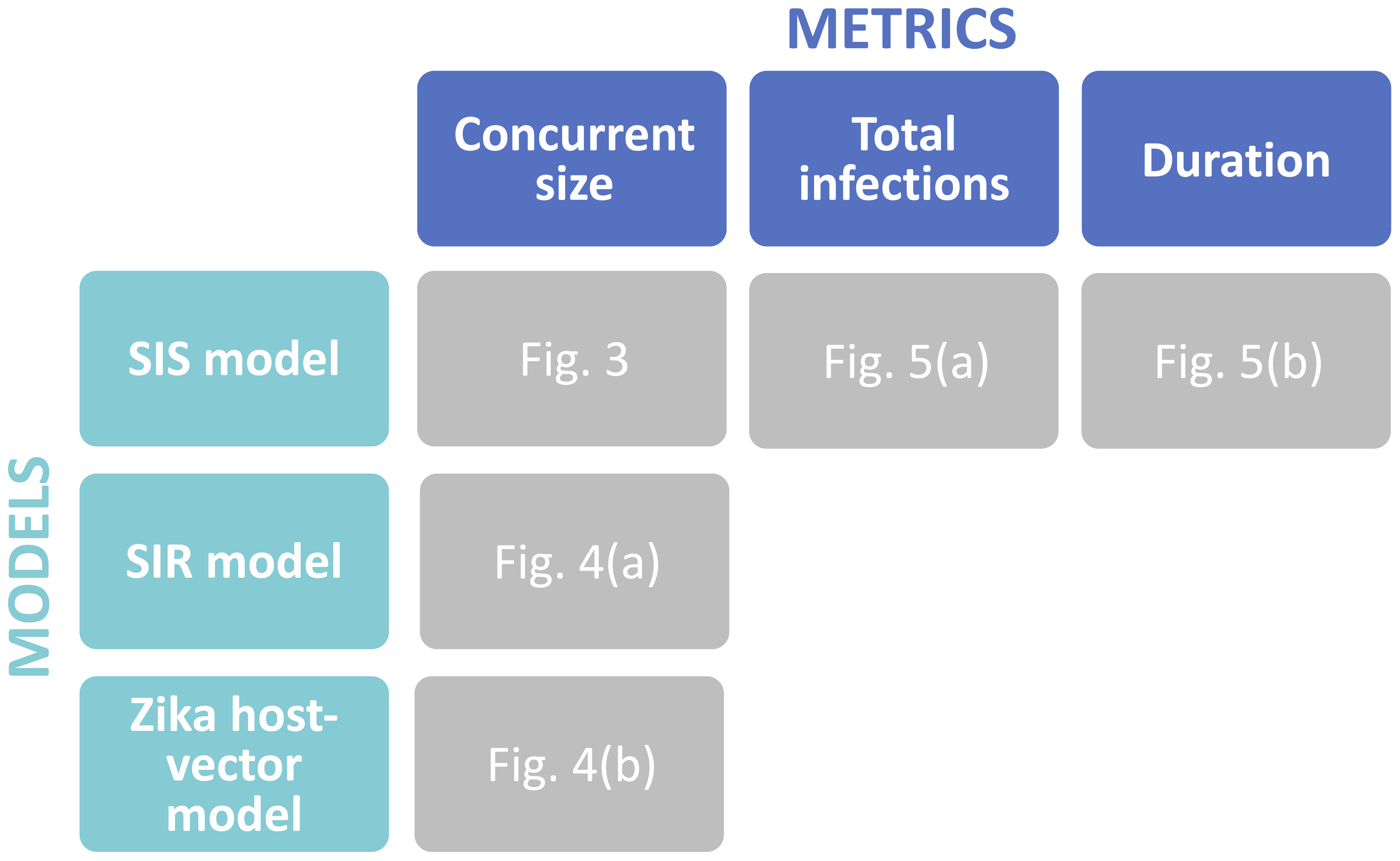
