## Supplementary material for "Will an outbreak exceed available resources for control? Estimating the risk from invading pathogens using practical definitions of a severe epidemic": Text S1

**AUTHORS**

R.N. Thompson^1,2,^*, C.A. Gilligan^3^, N.J. Cunniffe^3^

**AFFILIATIONS**

^1^Mathematical Institute, University of Oxford, Oxford, UK

^2^Christ Church, University of Oxford, Oxford, UK

^3^Department of Plant Sciences, University of Cambridge, Cambridge, UK

In this Supplementary Text, we first derive the branching process estimate for the probability of a major epidemic under the Susceptible-Infected-Susceptible (SIS) and Susceptible-Infected-Removed (SIR) models. We then derive the maximum number of infected individuals over the course of the outbreak for the deterministic SIS and SIR models.

Probability of a major epidemic (branching process estimate for stochastic SIS/SIR models)

To derive the branching process estimate for the probability of a major epidemic according to the stochastic SIS or SIR models, we denote by *q*_I_ the probability that a major epidemic does not occur, starting from *I* infected individuals (with the remainder of the population assumed susceptible). Conditioning on whether the first event is an infection or recovery/removal event gives

$$q_{I}=\frac{\beta IS}{\beta IS+\mu I}q_{I+1}+\frac{\mu I}{\beta IS+\mu I}q_{I-1},$$

for *I* = 1, 2, . . ., where *q*_0_ = 1 since a major epidemic certainly does not occur if there are initially 0 infected individuals. Dividing the top and bottom of each fractional term by the recovery/removal rate, $\mu$, and noting that early in an outbreak $S\approx N$, gives

$$q_{I}\approx\frac{R_{0}I}{R_{0}I+I}q_{I+1}+\frac{I}{R_{0}I+I}q_{I-1},$$

where $R_{0}=\frac{\beta N}{\mu}$.

For a major epidemic to fail to occur starting from two infected individuals, it is necessary for each infection lineage to fade out. Assuming that lineages from both infected individuals are approximately independent gives $q_{2}\approx q_{1}^{2}$. This leads to an equation for the probability of a major epidemic occurring, starting from one infected individual, $q_{1}$, given by

$$q_{1}\approx\frac{R_{0}}{R_{0}+1}q_{1}^{2}+\frac{1}{R_{0}+1}.$$

This quadratic equation has solutions *q*_1_ = 1 and *q*_1_ = 1/*R*_0_. Markov chain theory states that the probability that a major epidemic does not occur is given by the minimal solution to this equation [1], so that the probability of a major epidemic is approximately equal to

$$\mathrm{Prob}\left( major epidemic \right)=1-q_{1}\approx\left\{ \begin{aligned} 0 \mathrm{for} R_{0}\leq1, \\ 1-\frac{1}{R_{0}} \mathrm{for} R_{0}>1, \end{aligned} \right.$$

i.e. the expression given in the Methods section of the main text.

Maximum number simultaneously infected (deterministic SIS model)

The deterministic SIS model, given by system of equations (2) in the main text, can be solved explicitly for *I*(t) by replacing *S* in the second equation by *N – I*(t), separating variables and integrating, giving

$$I\left( t \right)=N\left( 1-\frac{1}{R_{0}} \right)\left( \frac{1}{A\exp\left( -\mu\left( R_{0}-1 \right)t \right)+1} \right).$$

The maximum number of infected individuals is then

$$I_{\max}=\left\{ \begin{aligned} N\left( 1-\frac{1}{R_{0}} \right) \mathrm{for} I\left( 0 \right)\leq N\left( 1-\frac{1}{R_{0}} \right), \\ I\left( 0 \right) \mathrm{for} I\left( 0 \right)>N\left( 1-\frac{1}{R_{0}} \right). \end{aligned} \right.$$

Maximum number simultaneously infected (deterministic SIR model)

In the deterministic SIR model (system of equations (3) in the main text), we first assume that $I(0)<I_{\max}$. In that case, the maximum value of $I(t)$ must occur when $\frac{dI}{dt}=0$, so that $S=\frac{\mu}{\beta}$. Dividing the second equation of system of equations (3) in the main text by the first, and then solving for *I*(t) in terms of *S*(t) gives

$$I\left( t \right)=-S(t)+\frac{\mu}{\beta}\ln\left( S(t) \right)+S\left( 0 \right)+I\left( 0 \right)-\frac{\mu}{\beta}\ln\left( S(0) \right).$$

At the maximum value of *I*(t), when $S=\frac{\mu}{\beta}$, we have

$$I_{\max}=-\frac{\mu}{\beta}+\frac{\mu}{\beta}\ln\left( \frac{\mu}{\beta} \right)+S\left( 0 \right)+I\left( 0 \right)-\frac{\mu}{\beta}\ln\left( S\left( 0 \right) \right),$$

which can be rearranged to give

$$I_{\max}=-\frac{N}{R_{0}}+\frac{N}{R_{0}}\ln\left( \frac{N}{{S\left( 0 \right)R}_{0}} \right)+S\left( 0 \right)+I\left( 0 \right).$$

Considering also the scenario in which *I*(*0*) is larger than this value, in which case the largest value of *I*(t) is *I*(0), gives

$$I_{\max}=\left\{ \begin{aligned} -\frac{N}{R_{0}}+\frac{N}{R_{0}}\ln\left( \frac{N}{{S\left( 0 \right)R}_{0}} \right)+S\left( 0 \right)+I\left( 0 \right) \mathrm{for} I\left( 0 \right)\leq-\frac{N}{R_{0}}+\frac{N}{R_{0}}\ln\left( \frac{N}{{S\left( 0 \right)R}_{0}} \right)+S\left( 0 \right)+I\left( 0 \right), \\ I\left( 0 \right) \mathrm{for} I\left( 0 \right)>-\frac{N}{R_{0}}+\frac{N}{R_{0}}\ln\left( \frac{N}{{S\left( 0 \right)R}_{0}} \right)+S\left( 0 \right)+I\left( 0 \right). \end{aligned} \right.$$
