## Supplementary material for "Will an outbreak exceed available resources for control? Estimating the risk from invading pathogens using practical definitions of a severe epidemic": Text S2

**AUTHORS**

R.N. Thompson^1,2,^*, C.A. Gilligan^3^, N.J. Cunniffe^3^

**AFFILIATIONS**

^1^Mathematical Institute, University of Oxford, Oxford, UK

^2^Christ Church, University of Oxford, Oxford, UK

^3^Department of Plant Sciences, University of Cambridge, Cambridge, UK

In the main text, we use simple models to demonstrate how the probability of a severe epidemic can be calculated for different definitions of a severe epidemic. One such definition uses the “concurrent size” metric, under which a severe epidemic is classified as an outbreak in which the number of individuals infected simultaneously exceeds the capacity for treatment. In this Supplementary Text, we show how the probability of a severe epidemic can be estimated under the “concurrent size” definition using simulations of three different models of an Ebola virus disease epidemic with varying levels of complexity.

Initially, we consider the Susceptible-Exposed-Infectious-Removed (SEIR) model and estimate the probability that the number of individuals infected simultaneously exceeds a threshold value. Then, we extend this to consider a more complex model in which Ebola Treatment Units (ETUs) are included explicitly and estimate the probability that the number of individuals simultaneously requiring ETU beds exceeds a threshold value. Finally, we extend this to consider a model in which ETUs are included and in which some cases do not report disease and therefore do not attend ETUs. This analysis demonstrates how the basic analyses in the main text can be extended for models that include additional features of real-world outbreaks.

Ebola virus disease outbreak models

A range of models were used during the 2014-16 Ebola epidemic in West Africa [1] to assess different possible control interventions. Among the simplest models used was the SEIR model; the deterministic SEIR model is given by the system of equations

$$\frac{dS}{dt}=-\beta IS,$$

$$\frac{dE}{dt}=\beta IS-\gamma E,$$

$$\frac{dI}{dt}=\gamma E-\mu I,$$

$$\frac{dR}{dt}=\mu I.$$

This model is an extension of the SIR model (system of equations (3) in the main text), but with individuals passing through a latent period (represented by the *E* compartment) with mean $1/\mu$ days. In the SEIR model, $R_{0}=\frac{\beta N}{\mu}$.

An important elaboration to basic compartmental epidemic models during the 2014-16 Ebola epidemic in West Africa was the inclusion of individuals in ETUs (e.g. [2]). Infected individuals in ETUs (*T*) can be included in the SEIR model, giving the following model (the SEITR model):

$$\frac{dS}{dt}=-\beta IS,$$

$$\frac{dE}{dt}=\beta IS-\gamma E,$$

$$\frac{dI}{dt}=\gamma E-\mu I,$$

$$\frac{dT}{dt}=\mu I-\theta T,$$

$$\frac{dR}{dt}=\theta T.$$

In the SEITR model, $R_{0}=\frac{\beta N}{\mu}$.

The third model that we consider here is an extension of the SEITR model to include individuals that do not report disease (*U*), giving the SEITUR model:

$$\frac{dS}{dt}=-\beta I(S+U),$$

$$\frac{dE}{dt}=\beta I(S+U)-\gamma E,$$

$$\frac{dI}{dt}=r\gamma E-\mu_{1}I,$$

$$\frac{dT}{dt}=\mu_{1}I-\theta T,$$

$$\frac{dU}{dt}=\left( 1-r \right)\gamma E-\mu_{2}U,$$

$$\frac{dR}{dt}=\theta T+\mu_{2}U.$$

In this model, individuals leaving the *E* class either enter the *I* class (with ascertainment probability $r$), after which they report disease and are transferred to an ETU, or do not report disease (with probability $1 - r$) and enter the *U* class (which represents unascertained cases). In the SEITUR model, $R_{0}=r\frac{\beta N}{\mu_{1}}+(1-r)\frac{\beta N}{\mu_{2}}$.

Parameters used to simulate the models in this supplementary analysis are shown in Tables S1-S3. We run simulations of the analogous stochastic models to the three models above using the Gillespie direct method [3].

Probability of a severe epidemic (“concurrent size” metric)

Motivated by the use of epidemic models to predict the numbers of ETU beds required in different locations during the 2014-16 Ebola epidemic in West Africa, we ran simulations of the stochastic SEIR, SEITR and SEITUR models to estimate the probability of a severe epidemic according to the “concurrent size” definition of a severe epidemic (Fig S3). In the SEIR model, this corresponded to the value of *I* exceeding a pre-specified threshold value at any point during the outbreak. In the SEITR and SEITUR models, we instead considered whether or not the number of individuals requiring treatment in ETUs (*T*) ever exceeds the threshold value.

For the parameters considered here, the SEIR and SEITR models generated similar estimates of the probability of a severe epidemic (Fig S3a,b). The required number of treatment beds would be estimated to be slightly lower using the SEITR model than using the SEIR model, since infectious individuals only require an ETU bed in the SEITR model when they enter the *T* class (i.e. not immediately on leaving the *E* class, unlike in the SEIR model).

However, when some cases were assumed not to report disease and the SEITUR model was used, the predicted number of ETU beds required, as well as the probability of a severe epidemic, were substantially different. First, for a fixed value of $R_{0}$, fewer beds were estimated to be required than under the SEIR and SETIR models, since not all individuals report to an ETU. Second, when $R_{0}$ was larger than but not close to one, the “plateau” (which corresponds to the branching process estimate for the probability of a major epidemic) was lower for the SEITUR model compared to the SEIR and SEITR models. We note that, in the SEITUR model, not only are unascertained cases included, but the infectious period distribution is no longer exponential (since an infected individual has an infectious period drawn from an exponential distribution with mean ${1/\mu}_{1}$ with probability *r*, and from a different exponential distribution with mean ${1/\mu}_{2}$ with probability 1 – *r*).

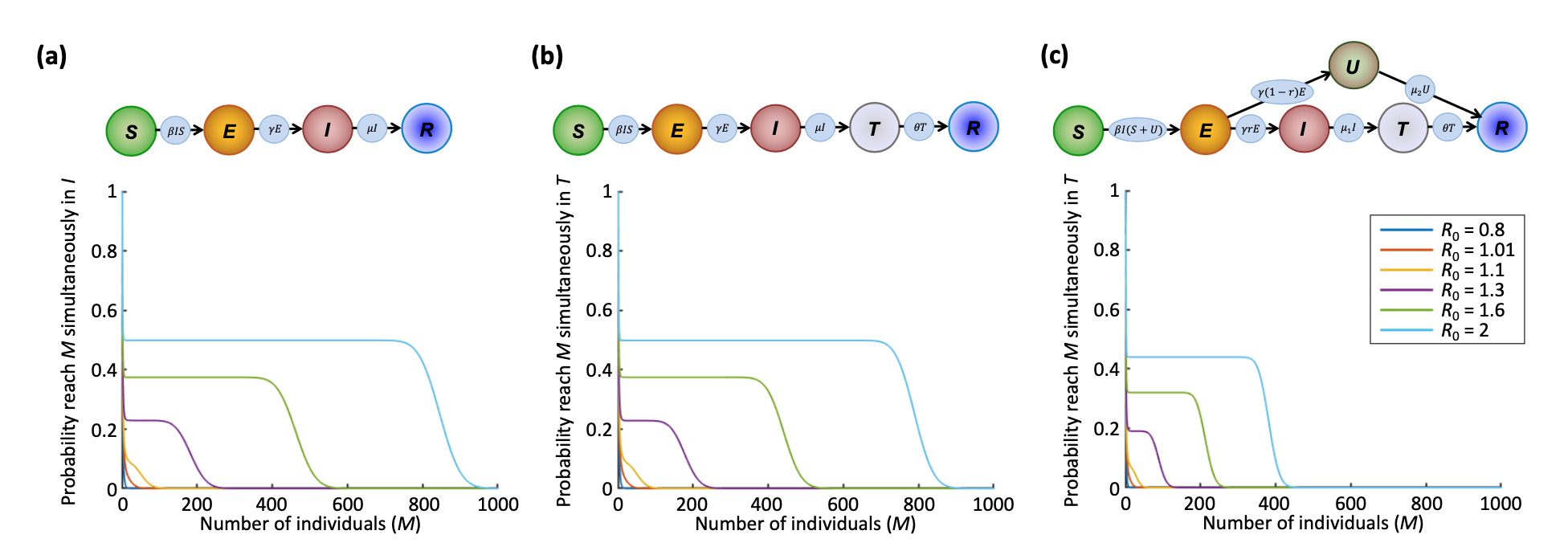

Figure S3. Probability of a severe epidemic for different models of an Ebola outbreak, where a severe epidemic is defined as an outbreak in which *M* simultaneously infected individuals is exceeded (“concurrent size” metric). (a) SEIR model. (b) SEITR model. (c) SEITUR model. In each panel, $R_{0}$ is varied by changing the value of $\beta$. 1,000,000 simulations were run for each value of *M* to estimate the probability of a severe epidemic. Each simulation was run in a population of *N* = 10,000 individuals, with initial conditions *E*(0) = 1 and the remaining *N* – 1 individuals assumed to be susceptible. Other model parameters are shown in Tables S1 (panel a), S2 (panel b) and S3 (panel c).

| **Parameter** | **Meaning** | **Value** |
| --- | --- | --- |
| $\beta$ | Infection rate | Varied to set the value of $R_{0}$ |
| $1/\gamma$ | Latent period | 9.4 days |
| $1/\mu$ | Infectious period | 10.9 days |

Table S1. Parameter values used in simulations of the stochastic SEIR model [2].

| **Parameter** | **Meaning** | **Value** |
| --- | --- | --- |
| $\beta$ | Infection rate | Varied to set the value of $R_{0}$ |
| $1/\gamma$ | Latent period | 9.4 days |
| $1/\mu$ | Infectious period (period until admitted to ETU) | 4 days |
| $1/\theta$ | Period in ETU | 6.9 days |

Table S2. Parameter values used in simulations of the stochastic SEITR model [2].

| **Parameter** | **Meaning** | **Value** |
| --- | --- | --- |
| $\beta$ | Infection rate | Varied to set the value of $R_{0}$ |
| $1/\gamma$ | Latent period | 9.4 days |
| $r$ | Ascertainment (reporting) probability | 0.6 |
| $1/\mu_{1}$ | Infectious period (period until admitted to ETU) | 4 days |
| $1/\theta$ | Period in ETU | 6.9 days |
| $1/\mu_{2}$ | Infectious period for unascertained cases | 10.9 days |

Table S3. Parameter values used in simulations of the stochastic SEITUR model [2].
